# DuckSeg: rapid, low-intervention assessment of protein recruitment after laser microirradiation

**DOI:** 10.64898/2026.11.27.769642

**Authors:** Istvan Kato, Matyas Susits, Istvan Szepesi-Nagy, Gergely Rona

## Abstract

**Motivation:** Laser microirradiation microscopy enables dynamic measurements of DNA repair protein recruitment, yet evaluating multiple recordings can require time-consuming image processing before an experimental overview is available. We developed a streamlined workflow to convert time-lapse recordings and irradiation masks into readily interpretable recruitment plots with minimal user intervention, facilitating early comparison of experimental conditions and informed selection of datasets for further analysis.

**Results:** DuckSeg integrates CellSAM nuclear segmentation, frame-to-frame mask association, fluorescence quantification within irradiation tracks relative to the surrounding nucleoplasm, and automated quality control in a Python-based, batch-oriented workflow. It provides condition-level recruitment curves based on complementary fluorescence difference and ratio scores, exportable single-cell measurements, and interactive Jupyter and 3D visualizations for optional rapid curation. Analysis of XRCC1, NEIL3, MSH2, MSH3, and EXO1b recordings demonstrates DuckSeg’s ability to recover established recruitment phenotypes across genetic and pharmacological perturbations. An additional NEIL3 dataset further illustrates the applicability of the workflow across different fluorescent reporter configurations. By combining automated processing with accessible visual inspection, DuckSeg supports efficient first-pass assessment of laser microirradiation experiments.

**Availability and implementation:** Open-source MIT-licensed code: https://github.com/ronalabrcns/DuckSeg; API documentation is available at https://duckseg.readthedocs.io.

## 1 Introduction

Laser microirradiation combined with live-cell fluorescence microscopy provides a spatially controlled means of monitoring DNA repair proteins following localized damage. Fluorescence accumulation or depletion within irradiated nuclear regions reveals the timing and persistence of repair responses (Miwatani-Minter and Rona, 2021; Rona, et al., 2018). However, extracting reliable recruitment curves from time-lapse recordings requires consistent identification of nuclei and irradiation regions, despite cell motion, variable signal intensity, and photobleaching (Bianchini and Kurz, 2023; Mahadevan, et al., 2019).

Several tools facilitate image-based analysis of protein dynamics. Fiji/ImageJ, CellProfiler, and TrackMate offer flexible image processing, segmentation, and tracking capabilities (Carpenter, et al., 2006; Ershov, et al., 2022; Schindelin, et al., 2012). Multistep ImageJ-based workflows have also been widely used in laser microirradiation studies, including quantification of fluorescence within irradiation stripes relative to nuclear reference regions (Miwatani-Minter and Rona, 2021; Rona, et al., 2024; Rona, et al., 2018). Dedicated tools include the Damage Analyzer Fiji scripts, which quantify background-corrected stripe enrichment from regions drawn by the user for each cell (Gaudreau-Lapierre, et al., 2018), and Stripenator, an ImageJ macro that detects nuclei and damage areas in multi-channel images using nuclear and damage-marker channels (Oeck, et al., 2019). Specialized software also includes CellTool, which integrates configurable image analysis, tracking, and mathematical fitting of recruitment kinetics, and qFADD.py, which couples drift-corrected fluorescence extraction with diffusion-based modeling (Bowerman, et al., 2022; Danovski, et al., 2023). These tools provide powerful options for detailed quantitative analysis, and DuckSeg is not intended to replace them.

Yet an important practical need arises earlier in the experimental process: rapidly determining whether proteins are recruited, whether experimental conditions produce discernible differences, and which datasets warrant more detailed examination. This is particularly useful when optimizing microscopy settings, screening perturbations, or reviewing large batches of recordings. An approach geared toward generating interpretable, condition-level recruitment curves with limited intervention would complement existing analytical tools.

To address this need, we developed DuckSeg (<u>D</u>eep-learning <u>U</u>tility for <u>C</u>ell mas<u>K</u>ing and Segmentation), an open-source tool designed to transform laser microirradiation recordings into interpretable protein recruitment curves with minimal user intervention. By enabling rapid assessment of recruitment dynamics across multiple experimental conditions, DuckSeg facilitates preliminary screening, experimental troubleshooting, and the identification of datasets warranting more detailed investigation. Here, we demonstrate its utility by analyzing published DNA repair datasets and additional microscopy recordings encompassing genetic perturbations, pharmacological treatments, and alternative fluorescent reporter configurations.

## 2 Implementation

DuckSeg is implemented as a modular Python-based workflow operated through interactive Jupyter notebooks (**Fig. 1, Supplementary Fig. S1**). Users provide time-lapse microscopy recordings and corresponding irradiation-region masks, which are processed in batches through sequential image-analysis and quantification steps. Intermediate results can be inspected through interactive visualization tools, allowing optional manual adjustments and quality control. Segmentation and tracking results are cached, so downstream steps can be repeated without re-running segmentation.

**Figure 1.**
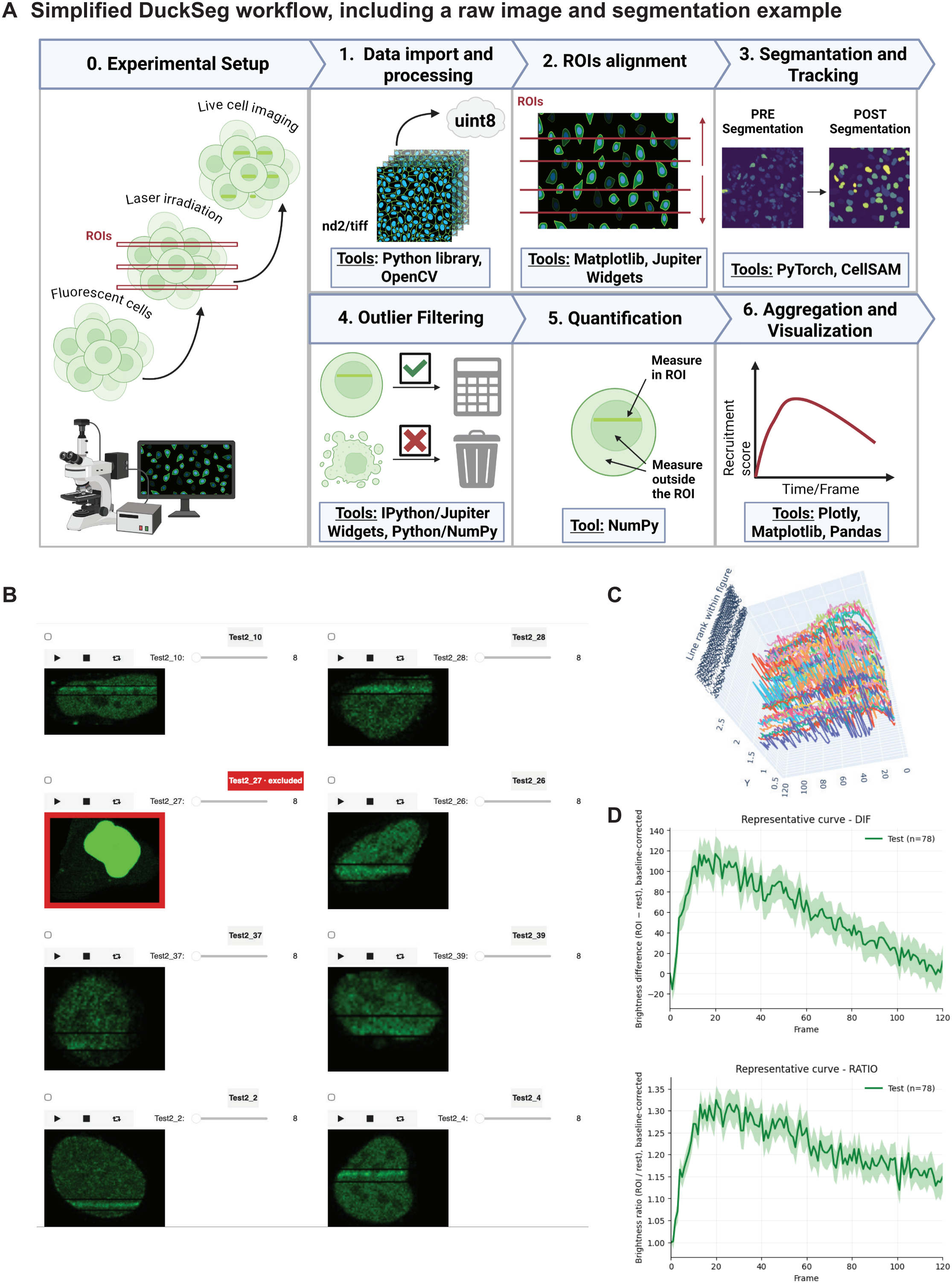
Overview of the DuckSeg image-analysis workflow. (A) Schematic overview of laser microirradiation microscopy, data import and preprocessing, ROI definition, CellSAM-based nuclear segmentation and temporal association, quality control, fluorescence quantification, and data visualization. (B) Interactive Jupyter-based viewer for inspecting individual-cell recordings and excluding unreliable trajectories. (C) Interactive 3D visualization of single-cell recruitment trajectories. (D) Representative automatically generated Diff- and Ratio-based recruitment curves. Panel A: Created in BioRender. Rona, G. (2027) https://BioRender.com/5pf3jao. *Alt text:* Schematic overview of the DuckSeg workflow, from laser microirradiation recordings to nuclear segmentation, tracking, quality control, and fluorescence quantification. Screenshots show the interactive cell-review interface, 3D visualization of single-cell recruitment trajectories, and automatically generated recruitment curves.

### 2.1 Data import, segmentation, and temporal linking

DuckSeg supports single-channel Nikon ND2 and multipage TIFF time-lapse recordings accompanied by binary masks defining the laser-irradiated regions of interest (ROIs). Recordings and their corresponding ROI masks are automatically identified using standardized file-naming conventions and organized into experimental batches. When appropriate, recordings can be restricted to a common number of frames to facilitate comparison across experimental conditions.

Images undergo configurable preprocessing for segmentation, including conversion to 8-bit, mean-brightness normalization, and thresholding. These transformations are applied only to the segmentation input, while the original fluorescence intensity data are preserved for subsequent quantification. Nuclei are segmented from the reporter signal itself, without a separate nuclear marker. Nuclear boundaries are identified independently in each frame using CellSAM, a deep learning-based instance segmentation model (Marks, et al., 2025). Nuclei whose bounding boxes overlap the row range of an irradiation band in the initial frame are selected.

To associate individual nuclei across time, DuckSeg compares binary nuclear masks between consecutive frames and selects the detection whose mask differs from the current mask in the fewest pixels, provided the difference does not exceed 60% of the current mask area. Otherwise, the most recent valid mask is carried forward. This approach provides an automated means of assembling single-cell time-lapse trajectories from successive nuclear segmentations.

### 2.2 Fluorescence quantification

For each identified nucleus, DuckSeg quantifies fluorescence enrichment within the irradiated region relative to the surrounding nucleoplasm. In the current implementation, irradiation tracks are defined as horizontal bands derived from the input ROI masks. The position and width of these bands can be adjusted to accommodate variations in irradiation geometry and the spatial extent of the recruitment signal.

At each time point, the mean fluorescence intensity is calculated within the irradiation band (I_ROI) and the remaining nuclear area (I_ref). Two complementary recruitment scores are then determined: Diff=(I_ROI)-(I_ref); Ratio=(I_ROI)/(I_ref). By using the surrounding nucleoplasm as an internal reference, these metrics quantify local fluorescence enrichment without requiring a separate extracellular background measurement for initial exploratory analysis. Both scores are calculated for individual nuclei throughout the recording, generating time-resolved recruitment trajectories. For each cell, Diff scores are baseline-corrected by subtracting the value at the first, pre-irradiation time point, whereas Ratio scores are normalized by dividing by their value at that time point. Consequently, Diff and Ratio trajectories start at 0 and 1, respectively.

### 2.3 Automated and interactive quality control

To facilitate rapid identification of unreliable measurements, DuckSeg incorporates both automated and user-guided quality-control procedures. Morphological filters flag nuclei with implausible areas, abrupt changes in nuclear size between consecutive frames, or excessive variation in nuclear area over time.

An interactive Jupyter-based viewer enables synchronized inspection of individual-cell microscopy sequences and manual exclusion of incorrectly segmented or tracked nuclei (**Fig. 1B**). Complementary interactive 3D plots visualize single-cell recruitment trajectories, facilitating the identification of cellular heterogeneity and anomalous profiles (**Fig. 1C**). Automatically and manually identified outliers are combined into a unified exclusion list for subsequent analysis.

### 2.4 Data visualization and export

Following automated or optional manual quality control, DuckSeg aggregates the retained single-cell trajectories according to experimental condition and generates recruitment curves for both Diff and Ratio scores (**Fig. 1D**). These plots display mean fluorescence enrichment over time, with shaded regions indicating the standard error of the mean (SEM), providing an immediate graphical overview of protein recruitment dynamics and differences between experimental conditions. Cells with an undefined score in any frame, for example when the widened band misses the nucleus or covers it entirely, are excluded from these curves; for the figures in this manuscript, partially complete trajectories were retained (Supplementary Methods). The workflow can export baseline-corrected recruitment plots as image files, together with a CSV file holding, for every cell and frame, the mean intensities I_ROI and I_ref and the Diff and Ratio scores, either raw or baseline-corrected. These data can be further processed in external software for additional visualization, statistical analysis, or detailed kinetic characterization.

Detailed implementation parameters and analysis examples are provided in the Supplementary Methods and online documentation. Dataset-specific analysis parameters are listed in **Supplementary Table S1**.

### 2.5 Experimental datasets

Published recordings were obtained from experiments reported by (Rona, et al., 2024) and (Miwatani-Minter and Rona, 2021). Additional N-terminally tagged mNeonGreen-NEIL3 recordings were acquired under the same experimental conditions as the published C-terminally tagged NEIL3 dataset (Supplementary Methods).

## 3 Applications

### 3.1 Assessment of XRCC1 and NEIL3 recruitment following D-type cyclin depletion

We first reanalyzed XRCC1 recruitment recordings comparing non-targeting siRNA controls (siNT) with cells depleted of all three D-type cyclins (siCyclin D1/D2/D3), previously investigated using a multistep ImageJ-based workflow, in which D-type cyclin depletion did not alter XRCC1 recruitment (Rona, et al., 2024). Fully automated DuckSeg analysis generated interpretable recruitment profiles, revealing post-irradiation fluorescence accumulation in both conditions (**Fig. 2A**). We compared these outputs with those obtained following optional manual quality control within DuckSeg, which allowed incorrectly segmented or tracked nuclei to be excluded. Manual curation excluded 68 of 145 trajectories (mostly because of lost tracks) and brought the curves of the two conditions closer together; the overall recruitment response was already apparent from the automated results. Moreover, additional manual filtering of the AUTO-derived dataset did not substantially alter the qualitative recruitment patterns (**Supplementary Fig. S2**). These findings illustrate DuckSeg’s ability to provide informative initial recruitment curves with limited user intervention.

**Figure 2.**
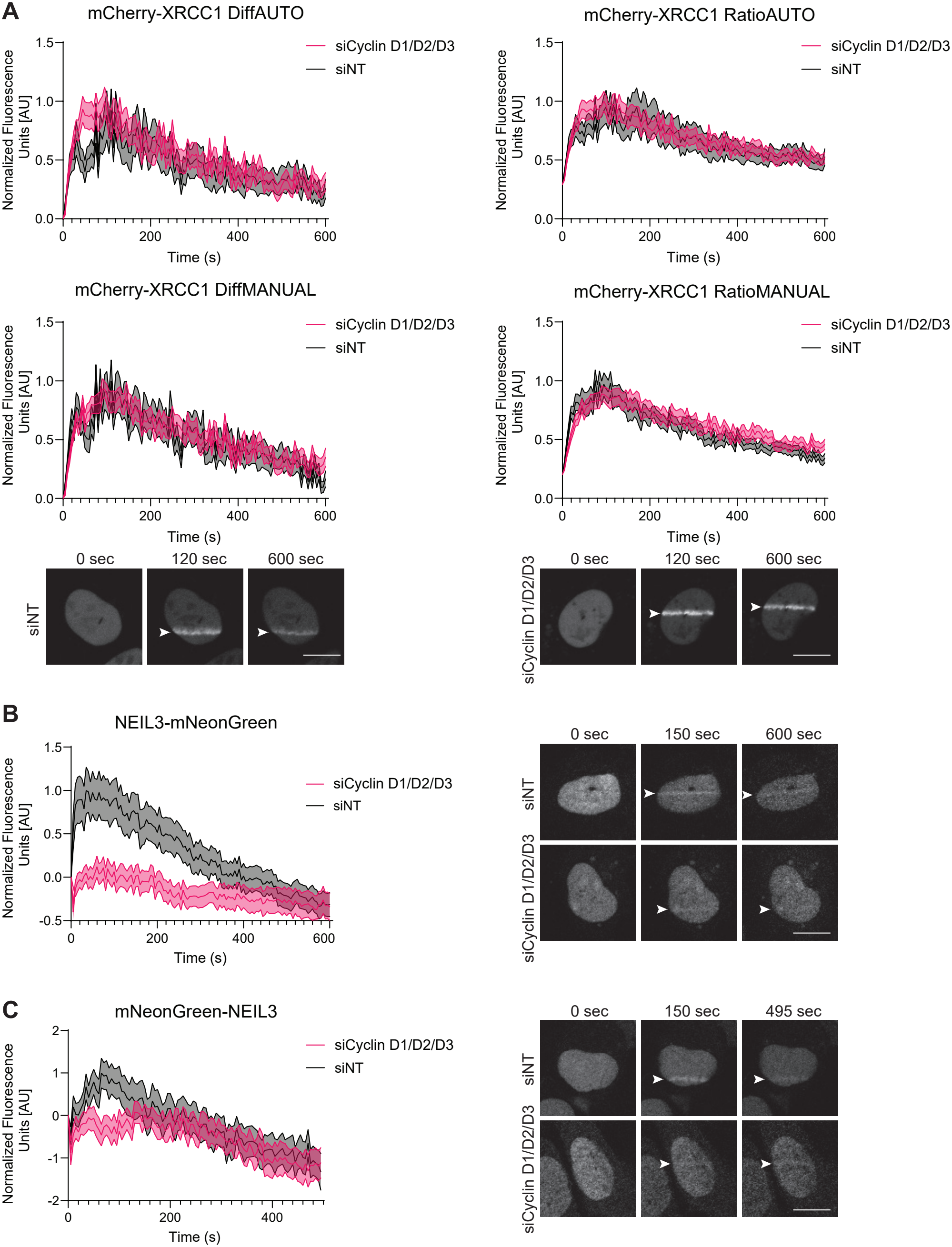
DuckSeg analysis of XRCC1 and NEIL3 recruitment following D-type cyclin depletion. (A) XRCC1 recruitment in non-targeting siRNA controls (siNT) and D-type cyclin-depleted cells (siCyclin D1/D2/D3), reanalyzed from previously published experiments (Rona, et al., 2024). Baseline-corrected fluorescence difference (Diff) and ratio (Ratio) scores are shown following fully automated processing (AUTO) and optional manual quality control within DuckSeg (MANUAL), together with representative time-lapse microscopy images. (B) Diff-based recruitment curves obtained by automated DuckSeg analysis (AUTO) and representative microscopy images for C-terminally tagged NEIL3-mNeonGreen, reanalyzed from previously published experiments (Rona, et al., 2024). (C) Diff-based recruitment curves from an independent dataset using N-terminally tagged mNeonGreen-NEIL3 following interactive quality control within DuckSeg (MANUAL), together with representative microscopy images. Curves represent mean ± SEM and were additionally normalized by setting the highest mean recruitment value within each graph to 1. Scale bars represent 10 μm. *Alt text:* Recruitment curves and representative fluorescence microscopy images of XRCC1 and two differently tagged NEIL3 reporters following D-type cyclin depletion. XRCC1 recruitment is shown following automated and manually curated analysis, while both NEIL3 reporters show reduced early recruitment following cyclin depletion.

To illustrate computational performance, DuckSeg was evaluated on five XRCC1 time-lapse recordings (121 frames each), containing 211 detected nuclei, of which 121 were selected for tracking based on their overlap with the irradiation bands. Segmentation and automated outlier filtering settings were identical to those used for the AUTO analysis in **Fig. 2A**. The combined segmentation and tracking run was completed in 17 min 18 s using Google Colab, demonstrating a practical processing time for obtaining an initial overview of multiple recordings (Supplementary Methods).

We next examined NEIL3 recruitment, previously shown to decrease following D-type cyclin depletion (Rona, et al., 2024). Reanalysis of the published NEIL3-mNeonGreen recordings reproduced this pronounced reduction in recruitment in cyclin-depleted cells (**Fig. 2B**). To further assess the applicability of the workflow, we analyzed an independent dataset using N-terminally tagged mNeonGreen-NEIL3. Despite greater variability and differences in the later recruitment profiles, this additional dataset also showed reduced early NEIL3 accumulation following D-type cyclin depletion (**Fig. 2C**). Together, these curves are qualitatively consistent with established recruitment phenotypes across independent datasets and alternative fluorescent reporter configurations.

### 3.2 Detection of recruitment responses across genetic and pharmacological perturbations

To further demonstrate the applicability of DuckSeg, we reanalyzed MSH2 and MSH3 recruitment datasets from cyclin D1 depletion-and-rescue experiments (Rona, et al., 2024). In the original study, depletion of D-type cyclins destabilized p21, reducing its competition with mismatch repair (MMR) proteins for proliferating cell nuclear antigen (PCNA) and thereby enhancing their recruitment to DNA lesions. This phenotype was rescued by expressing siRNA-resistant wild-type cyclin D1, but not a mutant defective in p21 binding and stabilization. The experiments compared empty-vector controls treated with non-targeting siRNA (EV_siNT), cyclin D1-depleted cells (EV_siUTR), and cells expressing wild-type (D1(WT)_siUTR) or p21-binding-defective cyclin D1 (D1(HP)_siUTR). DuckSeg recapitulated the increased recruitment of MSH2 and MSH3 upon cyclin D1 depletion and its suppression by wild-type, but not mutant, cyclin D1 (**Fig. 3A, B**).

**Figure 3.**
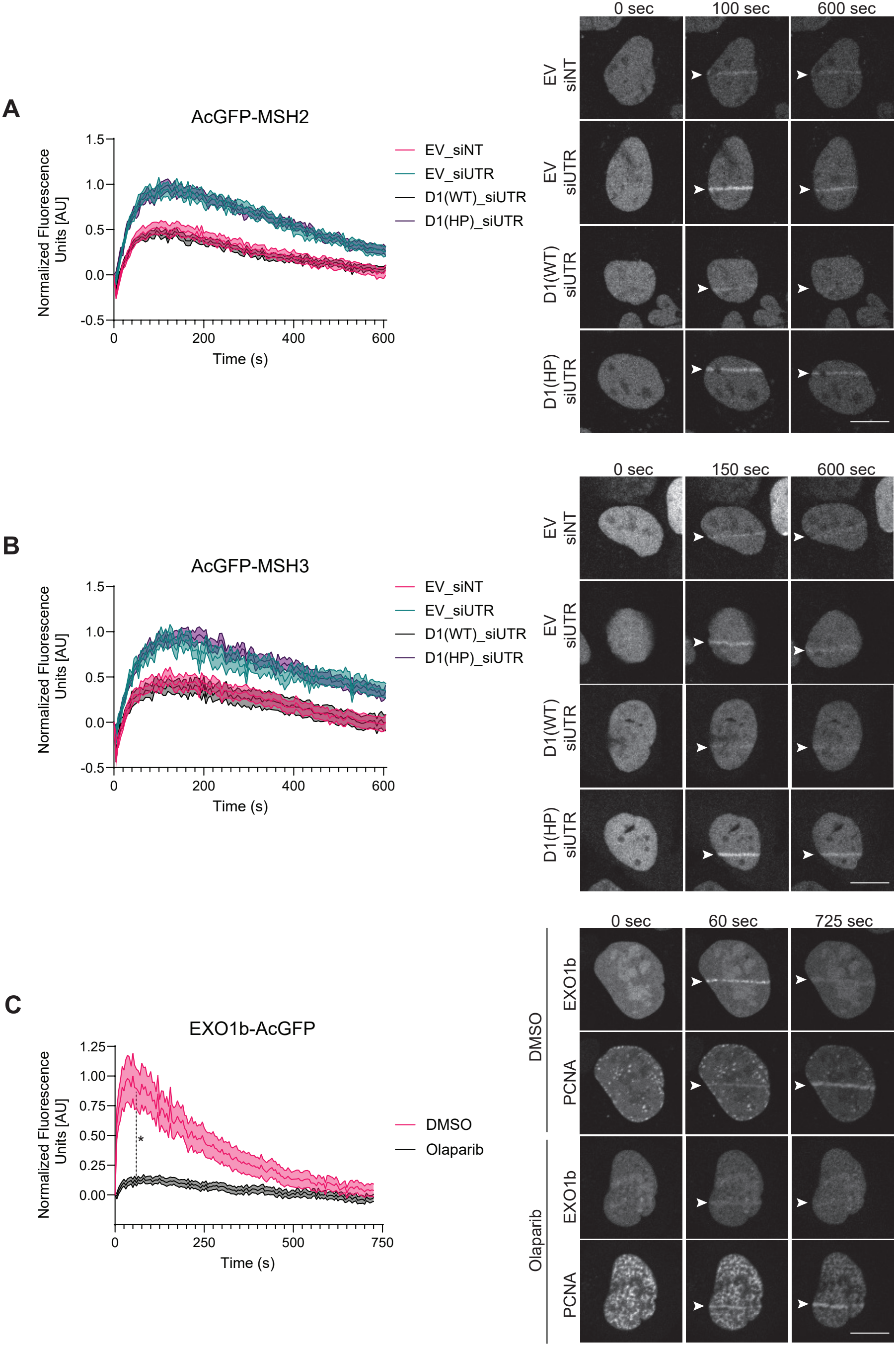
DuckSeg recapitulates DNA repair protein recruitment phenotypes following genetic and pharmacological perturbations. Recruitment kinetics quantified using baseline-corrected fluorescence difference (Diff) scores. (A,B) MSH2 and MSH3 recruitment in previously published cyclin D1 depletion-and-rescue experiments (Rona, et al., 2024). Conditions include empty-vector controls (EV_siNT), cyclin D1 depletion (EV_siUTR), and rescue with wild-type (D1(WT)_siUTR) or p21-binding-defective cyclin D1 (D1(HP)_siUTR). Representative time-lapse microscopy images illustrate recruitment under the four conditions. (C) EXO1b recruitment in cells treated with DMSO or the PARP1/2 inhibitor olaparib, reanalyzed from previously published experiments (Miwatani-Minter and Rona, 2021), restricted to manually selected S-phase nuclei, with corresponding EXO1b and PCNA fluorescence microscopy images. Olaparib significantly reduced recruitment at 60 seconds after irradiation (p < 0.0001, two-sided Mann–Whitney U test; indicated by an asterisk). Curves represent mean ± SEM and were additionally normalized by setting the highest mean recruitment value within each graph to 1. Scale bars represent 10 μm. *Alt text:* Recruitment curves and representative fluorescence microscopy images showing increased MSH2 and MSH3 recruitment following cyclin D1 depletion, reversed by wild-type but not mutant cyclin D1. EXO1b recruitment is reduced following olaparib treatment, with accompanying PCNA images illustrating S-phase cell selection.

As an independent pharmacological example, we reanalyzed EXO1b recruitment recordings from previously published S-phase laser microirradiation experiments (Miwatani-Minter and Rona, 2021). Because DuckSeg analyzes a single channel, S phase nuclei were identified by their punctate pattern in the PCNA channel, and all other tracked nuclei were removed through DuckSeg’s manual exclusion list. In the original study, PARP1/2 inhibition with olaparib significantly reduced EXO1b accumulation at laser-induced DNA damage sites. Consistent with these findings, DuckSeg curves showed reduced recruitment in olaparib-treated cells compared with vehicle controls (**Fig. 3C**).

Together, these examples demonstrate qualitative agreement between DuckSeg-derived recruitment profiles and previously published results obtained using conventional ImageJ-based analysis across different DNA repair proteins, genetic perturbations, and pharmacological treatments.

## 4 Discussion

DuckSeg provides a streamlined approach for rapid evaluation of protein recruitment following laser microirradiation, bridging the gap between image acquisition and detailed quantitative analysis. By integrating automated image processing, fluorescence quantification, and interactive quality control, it enables researchers to assess recruitment dynamics across multiple experimental conditions with minimal user intervention.

Beyond retrospective analysis, DuckSeg could facilitate experimental screening and optimization. Rapid assessment of pilot recordings may help researchers refine irradiation conditions and imaging parameters during microscopy sessions (Zhao, et al., 2023). Interactive single-cell visualization and optional manual curation allow users to examine cellular heterogeneity, identify questionable trajectories, and refine recruitment profiles when necessary. The XRCC1 analysis illustrates that the overall recruitment response can be identified automatically, while optional manual curation can refine comparisons between experimental conditions. Thus, DuckSeg complements existing comprehensive analysis platforms by emphasizing rapid, accessible assessment of experimental outcomes.

Several limitations should be considered. Segmentation and frame-to-frame association can introduce errors, particularly when nuclei move, overlap, or change morphology. Although automated and manual quality control can identify problematic trajectories, some inaccuracies may remain. The current implementation analyzes single-channel recordings, assumes horizontal irradiation tracks spanning the field, and does not subtract extracellular background. The Ratio score is analogous to a previously described enrichment measure (Gaudreau-Lapierre, et al., 2018) and can compensate for uniform photobleaching, whereas the Diff score is insensitive to spatially uniform additive background but remains sensitive to bleaching. SEM bands are calculated from cells pooled across recordings and therefore reflect the precision of the pooled cell-level mean rather than variability between independent experiments (Lazic, 2010). DuckSeg is therefore primarily suited to exploratory analysis.

Overall, DuckSeg offers a practical means of accelerating the initial interpretation of laser microirradiation experiments and guiding subsequent experimental decisions.

## Availability and implementation

DuckSeg is freely available under the MIT license at https://github.com/ronalabrcns/DuckSeg, with documentation at https://duckseg.readthedocs.io/en/latest/. The version described here (v1.1.0) is archived at Zenodo (https://doi.org/10.5281/zenodo.24546970). The software is implemented as a modular Python package (Python ≥3.10), integrating CellSAM and PyTorch for deep learning-based nuclear segmentation, OpenCV for image processing, and standard scientific Python libraries for data analysis and visualization. DuckSeg operates through interactive Jupyter notebooks, with example workflows provided in the repository. The software supports local installation, with a Google Colab implementation provided to simplify access without dedicated local computational infrastructure. CellSAM model weights are obtained separately from Hugging Face under their own non-commercial license, which users must accept before first use.

## Data availability

The analyses include laser microirradiation recordings from previously published experiments (Miwatani-Minter and Rona, 2021; Rona, et al., 2024), and additional recordings of N-terminally tagged mNeonGreen-NEIL3. An example dataset with ROI masks, used by the Colab notebooks, is available from the DuckSeg GitHub repository (https://github.com/ronalabrcns/DuckSeg/releases/tag/example-data). The recruitment summary data underlying Figures 2 and 3 and Supplementary Figure S2 are provided in Supplementary Table S2. The raw recordings are available from the corresponding author upon request.

## Author contributions

I.K., M.S., and I.S.-N. designed, implemented, and tested the software and contributed to its development. I.K. and I.S.-N. prepared code documentation. I.K. conducted image reanalysis. I.K. and G.R. performed the additional NEIL3 imaging experiments. G.R. conceived and supervised the project. All authors contributed to writing and editing the manuscript.

## Acknowledgements

ChatGPT (OpenAI) was used to assist with language editing and text refinement. All content was reviewed and approved by the authors, who take full responsibility for the final manuscript.

