## Supplementary for "DuckSeg: rapid, low-intervention assessment of protein recruitment after laser microirradiation"

Running title: DuckSeg: rapid protein recruitment analysis

**Outline**

**Supplementary Methods**

**Supplementary Figures:**

*Figure S1 related to Figure 1 and Supplementary Methods*
*Figure S2 related to Figure 2*

**Supplementary Tables:**

*Table S1 related to Figures 2, 3 and S2*
*Table S2 related to Figures 2, 3 and S2*

### SUPPLEMENTARY METHODS

#### Laser microirradiation and live-cell fluorescence microscopy

The XRCC1, C-terminally tagged NEIL3-mNeonGreen, MSH2, and MSH3 datasets derive from previously published laser microirradiation experiments (Rona, et al., 2024). The EXO1b recordings derive from previously described experiments (Miwatani-Minter and Rona, 2021). The original time-lapse microscopy recordings and corresponding irradiation-region masks were reanalyzed using DuckSeg. For the EXO1b dataset, all stripe-crossing nuclei were tracked by DuckSeg, and nuclei with homogeneous PCNA signal (G1, G2 or M) were then excluded by cell identifier through the manual exclusion list, so that only nuclei with a punctate PCNA pattern (S phase) contributed to the curves.

An additional dataset was acquired using N-terminally tagged mNeonGreen-NEIL3. Cell line, cell preparation, synchronization, siRNA transfection, live-cell imaging conditions, and irradiation protocol were the same as those employed for the C-terminally tagged NEIL3 reporter in (Rona, et al., 2024), except for the orientation of the fluorescent fusion. Briefly, G1-synchronized U2OS cells stably expressing the reporter were compared after treatment with non-targeting siRNA (siNT) or siRNAs targeting CCND1, CCND2, and CCND3 (siCyclin D1/D2/D3). Repair-factor recruitment experiments were done on a Nikon A1R-HD25 confocal microscope equipped with a 60×/1.42-NA oil-immersion Plan-Apochromat objective. Localized irradiation employed a Nikon LUN-F 50-mW 405-nm FRAP laser unit at 100% power output and a 1,000-μs dwell time. Representative images in the main figures are selected from the corresponding analyzed recordings.

#### Input data, condition assignment, and frame handling

DuckSeg is a Python-based pipeline that integrates four core computational tasks for analyzing laser microirradiation datasets: (i) deep learning-based instance segmentation and temporal tracking of individual nuclei; (ii) quantification of fluorescence intensity within the irradiated region of interest (ROI) relative to the surrounding nucleoplasm; (iii) automated and interactive quality control; and (iv) aggregation and visualization of protein recruitment kinetics across experimental conditions.

Input data are organized into experimental batches, with each batch representing a defined experimental condition and containing time-lapse microscopy recordings in Nikon ND2 or multipage TIFF format, together with corresponding binary masks delineating the laser-irradiated ROIs. Recordings and masks are paired by file name: each mask shares the filename stem of its recording, followed by the suffix _ROI.tif. ROI masks must match the spatial dimensions of their recordings and identify the irradiated region through nonzero pixel values. Experimental conditions are assigned through batch organization, while individual recordings are identified by their filenames, eliminating the need for separate metadata manifests.

To facilitate comparisons across recordings, DuckSeg supports restricting all time-lapse sequences within a batch to a common number of frames. In the example workflow, the maximum sequence length is set to that of the shortest recording, ensuring that all analyzed trajectories cover an identical number of time points. This procedure truncates longer recordings without temporal interpolation or resampling. Because matching frame counts do not necessarily imply identical acquisition times, recordings should have consistent imaging intervals for direct comparison of recruitment kinetics.

For segmentation preprocessing, image data are converted to 8-bit. ND2 and 16-bit TIFF frames are scaled by 255/65,535 regardless of the detector's effective bit depth, whereas native 8-bit TIFF recordings are used unchanged. Importantly, the original fluorescence intensity data are preserved independently for subsequent quantitative analysis, ensuring that segmentation-specific preprocessing does not alter the underlying photometric measurements.

**Preprocessing, ROI alignment, and segmentation**

To optimize nuclear segmentation across recordings with variable fluorescence intensities and imaging conditions, DuckSeg incorporates a configurable image-preprocessing pipeline. Available transformations include conversion to 8-bit depth, mean-brightness normalization, linear contrast adjustment, and global intensity thresholding. These transformations are applied sequentially to the images used for segmentation, while the original fluorescence intensity data are preserved independently for subsequent quantitative analysis. Users can preview the effects of preprocessing on selected frames and adjust the parameters before processing the complete dataset.

In the documented example workflow, frames are converted to 8-bit representations, normalized to a target mean intensity of 20, and subsequently subjected to global binary thresholding at an intensity value of 20. Normalizing mean brightness before thresholding helps accommodate variations in fluorescence intensity between recordings and provides more consistent inputs for nuclear segmentation. However, the preprocessing sequence is fully configurable and can be adapted to different fluorophores, expression levels, and imaging conditions. The specific preprocessing, irradiation-band width, quality-control, and acquisition parameters used for each experimental dataset analyzed in this study are summarized in **Supplementary Table S1**.

To compensate for drift between frames, DuckSeg provides optional image-registration tools (ORB keypoint matching with homography estimation, and correlation-based alignment). These were not applied to the datasets analyzed here, as the subsequent frame-to-frame tracking procedure can accommodate modest nuclear displacement. Additionally, an interactive Jupyter-based ROI editor allows users to inspect the detected irradiation bands and manually adjust their positions when necessary. These corrections are incorporated into subsequent fluorescence quantification to ensure appropriate alignment between the irradiation regions and nuclear measurements.

Nuclear segmentation is performed independently in each frame using CellSAM (Marks, et al., 2025), a deep learning-based instance segmentation model (cellsam_general model, v1.2). The implementation utilizes CellSAM's default image normalization and applies a fixed (not user-configurable) bounding-box detection confidence threshold of 0.7. Following segmentation of the initial frame, DuckSeg selects nuclei for analysis whose detected bounding boxes overlap the row range of an irradiation band, treating these bands as spanning the full image width. CellSAM subsequently segments the remaining frames, and the resulting nuclear masks are linked across time to reconstruct individual-cell trajectories, as described in the following section.

#### Temporal association and tracking assumptions

To follow individual nuclei across frames, DuckSeg uses mask-based linking rather than bounding-box overlap alone, which can produce incorrect associations in dense fields. Each tracked nucleus is compared with every detection in the next frame by calculating the symmetric pixel-wise difference between masks in image coordinates, that is, the number of pixels belonging to only one of the two masks.

The detection with the smallest difference is selected if the difference does not exceed 60% of the area of the most recently assigned mask. For an approximately circular nucleus, this tolerates displacement of up to about half its radius between consecutive frames. If no detection meets this criterion, the most recent valid mask and bounding box are carried forward and applied to the current frame, and matching resumes when a suitable detection reappears. Trajectories are not terminated, so a nucleus that is permanently lost continues to be measured at its last valid position; these cases are not flagged automatically and are identified during quality control.

Matching is local and greedy, without global optimization or one-to-one assignment between tracks. Large movements, cell division, or abrupt morphological changes can therefore produce incorrect associations; a neighboring nucleus may be assigned to two trajectories, and partial or merged segmentations within the 60% tolerance may be linked. Such trajectories can be identified by automated morphological filtering and optional interactive review, as described below.

The resulting temporally associated nuclear masks, corresponding cropped fluorescence images, and intersections with the laser-irradiated regions provide the spatial information required for subsequent single-cell recruitment quantification.

#### Irradiation geometry and fluorescence scores

Laser-irradiated regions are defined using binary ROI masks specifying one or more approximately horizontal irradiation stripes. DuckSeg identifies these regions by scanning the central column of each ROI mask and extracting contiguous intervals of nonzero pixels, thereby defining the vertical positions and widths of the irradiation bands. Consequently, irradiation stripes must intersect the central image column to be detected. Each initially selected nucleus is associated with the irradiation band intersecting its bounding box. The band's row position is expressed relative to the upper edge of the nucleus's first-frame bounding box and applied at the same offset to the crop of every later frame. The band therefore follows vertical displacement of the bounding box and is sensitive to changes in the segmented upper boundary. In the current implementation, fluorescence quantification uses the first associated band when multiple irradiation regions are present.

To accommodate minor positional inaccuracies in ROI annotation and the spatial extent of protein accumulation around the irradiated region, DuckSeg allows the irradiation bands to be symmetrically expanded around their midpoints. The width of the quantification region is controlled by an adjustable multiplicative parameter (range_modifier), where a value of 1 preserves the original annotated width and larger values proportionally expand it (e.g., 5 increases the width fivefold). An interactive ROI editor additionally allows manual adjustment of irradiation-band positions when necessary, ensuring appropriate alignment with the observed recruitment signal.

For each tracked nucleus, fluorescence intensity is quantified separately within the expanded irradiation band and the remaining segmented nuclear area outside this region, which serves as an internal reference representing the surrounding nucleoplasm. Mean intensities are calculated using the original fluorescence measurements and normalized by the number of foreground nuclear pixels within each region, thereby accounting for differences in region size and nuclear morphology.

Let I_ROI(t) and I_ref(t) denote the mean fluorescence intensities within the irradiated region and the surrounding nucleoplasm, respectively, at time t. DuckSeg calculates two complementary recruitment scores:

$$\mathrm{Diff}(t) = I\_ROI(t) - I\_ref(t)$$

$$\mathrm{Ratio}(t) = I\_ROI(t) / I\_ref(t)$$

To account for differences in fluorescence distribution before irradiation, individual-cell trajectories are normalized to their initial values. For Diff, the initial score is subtracted from all subsequent measurements, whereas Ratio values are divided by their corresponding initial score:

$$\mathrm{Diff\_norm}(t) = \mathrm{Diff}(t) - \mathrm{Diff}(0)$$

$$\mathrm{Ratio\_norm}(t) = \mathrm{Ratio}(t) / \mathrm{Ratio}(0)$$

These transformations yield baseline values of 0 and 1 for Diff and Ratio, respectively, facilitating comparisons of recruitment dynamics between cells and experimental conditions. Both approaches use the surrounding nucleoplasm as an internal reference without requiring a separate extracellular background measurement. No dedicated photobleaching correction is performed.

**Automated exclusion, interactive review, and optional post-processing**

DuckSeg provides automated morphological filtering and optional interactive quality control to identify trajectories affected by segmentation artifacts, nuclear merging, or tracking instability. Three configurable area-based filters are available: (i) absolute nuclear area bounds, (ii) maximum area change between consecutive frames, and (iii) maximum overall area variation across the recording. Trajectories failing any selected filter are flagged for exclusion. Of the available automated morphological filters, only absolute nuclear-area filtering was used, where indicated in Supplementary Table S1.

More complex errors can be inspected in an interactive Jupyter-based interface showing synchronized time-lapse crops of individual tracked nuclei, where users can manually exclude cells. Automatically and manually excluded cells are combined into a single list, which is applied when calculating scores and condition-level curves. Manual exclusion was based on visual inspection of segmentation consistency, tracking continuity, and the identity of individual nuclei throughout the time-lapse recordings.

As a complementary tool, an interactive three-dimensional (3D) plot displays individual-cell recruitment trajectories, allowing users to assess heterogeneity and identify atypical profiles by their cell identifiers for further review. The plot represents single-cell kinetic measurements rather than three-dimensional nuclear segmentation or tracking.

To examine the influence of additional manual filtering, the automated XRCC1 dataset was reanalyzed after excluding trajectories with atypical recruitment profiles, including pronounced oscillations or recruitment values exceeding five times the maximum mean recruitment value. This post-processing was performed on exported measurements, independently of interactive curation within DuckSeg (**Supplementary Fig. S2**).

**Aggregation, graphical outputs, CSV export, and caching**

Following automated and optional manual quality control, retained single-cell trajectories are grouped according to their experimental conditions, and DuckSeg calculates the mean score and standard error of the mean (SEM) across cells at each time point. DuckSeg generates recruitment plots for both fluorescence difference (Diff) and ratio (Ratio) scores, displaying the mean response over time with shaded regions representing the SEM. These plots can be visualized directly within the Jupyter environment or exported as high-resolution image files at 300 dpi for subsequent analysis or presentation. The widened irradiation band is clamped to the boundaries of each nucleus crop. Trajectories containing undefined (NaN) or infinite values, which may arise when the band misses the nucleus or covers it entirely, or from division by zero, are excluded from the automatically generated DuckSeg curves, and the number of excluded cells is reported. For the manuscript figures, partially complete trajectories were retained in GraphPad Prism provided that a valid measurement was available at the initial time point, with only valid measurements contributing to the mean and SEM at each subsequent time point.

In addition to condition-level summaries, the workflow supports interactive visualization of individual-cell recruitment trajectories, allowing users to further examine cellular heterogeneity and identify unusual responses. DuckSeg plots and exports use frame indices; time in seconds was obtained by multiplying by the acquisition interval (5 s). For presentation in the manuscript figures, recruitment curves were further scaled in GraphPad Prism using the transformation Y = Y/K, where K was defined as the highest mean recruitment value across all experimental conditions and time points within each graph. The same scaling factor was applied to all conditions and their corresponding SEM values, setting the maximum mean recruitment value to 1 while preserving relative differences between conditions. Consequently, absolute recruitment amplitudes cannot be directly compared between independently scaled graphs. The mean, standard deviation (SD), and number of contributing cells (N) at each time point before this additional normalization are provided in **Supplementary Table S2**.

For downstream quantitative analysis, DuckSeg exports structured comma-separated value (CSV) files containing time-resolved fluorescence measurements for individual tracked cells. These include the mean fluorescence intensities within the irradiated band and surrounding nucleoplasm, together with their corresponding Diff and Ratio scores. When individual trajectories differ in length, shorter sequences are padded with empty values to maintain a consistent tabular structure. The export contains all cells passed to it, meaning any excluded cells must be removed beforehand. The exported measurements preserve the individual-cell information underlying the aggregated recruitment curves, enabling additional normalization, visualization, statistical comparisons, and detailed kinetic analyses using external software.

To improve computational efficiency, DuckSeg incorporates persistent disk caching of segmentation and tracking results for each recording. This allows users to modify quality control settings, irradiation-band width, and plots without repeating the computationally intensive CellSAM inference. Furthermore, if irradiation-band positions change, cells are automatically re-tracked using the stored segmentation. However, the cache does not record preprocessing settings; therefore, when preprocessing transformations affecting nuclear segmentation are modified, the complete segmentation workflow must be rerun without relying on the cache.

**Software environment, installation, and model access**

DuckSeg is an open-source Python package available under the MIT license at https://github.com/ronalabrcns/DuckSeg, with installation instructions and documentation at https://duckseg.readthedocs.io/. The software is organized into modular components for data import, image preprocessing, spatial registration, segmentation and tracking, interactive quality control, fluorescence quantification, and data export, allowing individual processing stages to be configured and inspected independently. The software can be executed through Jupyter notebooks on local computers or in Google Colab. Core dependencies include CellSAM for nuclear segmentation, PyTorch for deep learning inference, and standard Python libraries for image processing, visualization, and data analysis. GPU acceleration is recommended for computationally intensive segmentation tasks.

Computational performance was evaluated in Google Colab using a high-RAM Python 3 runtime equipped with an NVIDIA T4 GPU (51.0 GB system RAM, 15.0 GB GPU memory). Five XRCC1 time-lapse recordings, each containing 121 frames, were processed. The recordings contained 17, 37, 48, 51, and 58 initially detected nuclei, respectively. Of these, 14, 19, 25, 27, and 36 nuclei were selected for tracking based on the overlap of their initial bounding boxes with the irradiation bands. Individual segmentation and tracking times were 2 min 16 s, 2 min 29 s, 2 min 54 s, 3 min 18 s, and 3 min 37 s, respectively. Processing all five recordings in a single combined run required 17 min 18 s. These timings refer specifically to segmentation and tracking and exclude data preparation, interactive quality control, and downstream visualization.

CellSAM requires separately distributed pretrained model weights, which are subject to the model provider's licensing and access requirements. CellSAM model weights are obtained from the gated Hugging Face repository (vanvalenlab/cellsam). Users must accept the model license and provide a Hugging Face access token for the initial download. DuckSeg subsequently loads compatible cached weights without requiring another download. Alternatively, compatible model weights can be loaded from a local cache. A version-pinned dependency specification is provided in the GitHub repository to facilitate reproducible installation of the software environment. The DuckSeg source code is available at https://github.com/ronalabrcns/DuckSeg, and the version described here (v1.1.0) is archived at Zenodo (https://doi.org/10.5281/zenodo.24546970).

**Statistical analysis**

Recruitment curves in Figures 2, 3 and Supplementary Figure S2 display the arithmetic mean of valid single-cell measurements at each time point, with shaded bands indicating the standard error of the mean (SEM). Partially complete trajectories were retained for these figures, and the number of contributing cells (N) at each time point is reported in Supplementary Table S2. The curves in Figure 1D were generated directly by DuckSeg from complete trajectories only. These SEM bands describe the precision of the pooled cell-level mean and do not represent variability between independent experiments. Statistical comparisons between two experimental conditions were performed using the two-sided, nonparametric Mann-Whitney U test. For EXO1b, recruitment scores at 60 seconds after irradiation were compared between DMSO- and olaparib-treated cells ($p < 0.0001$). Statistical significance was defined as $p < 0.05$.

### SUPPLEMENTARY FIGURES

**Supplementary Figure S1. Step-by-step overview of the DuckSeg workflow.**
Steps from input recordings and irradiation masks through CellSAM segmentation, tracking, and quality control to Diff/Ratio scoring and export. Blue boxes indicate automated steps, orange boxes indicate user-interactive steps, and dashed borders indicate optional steps.

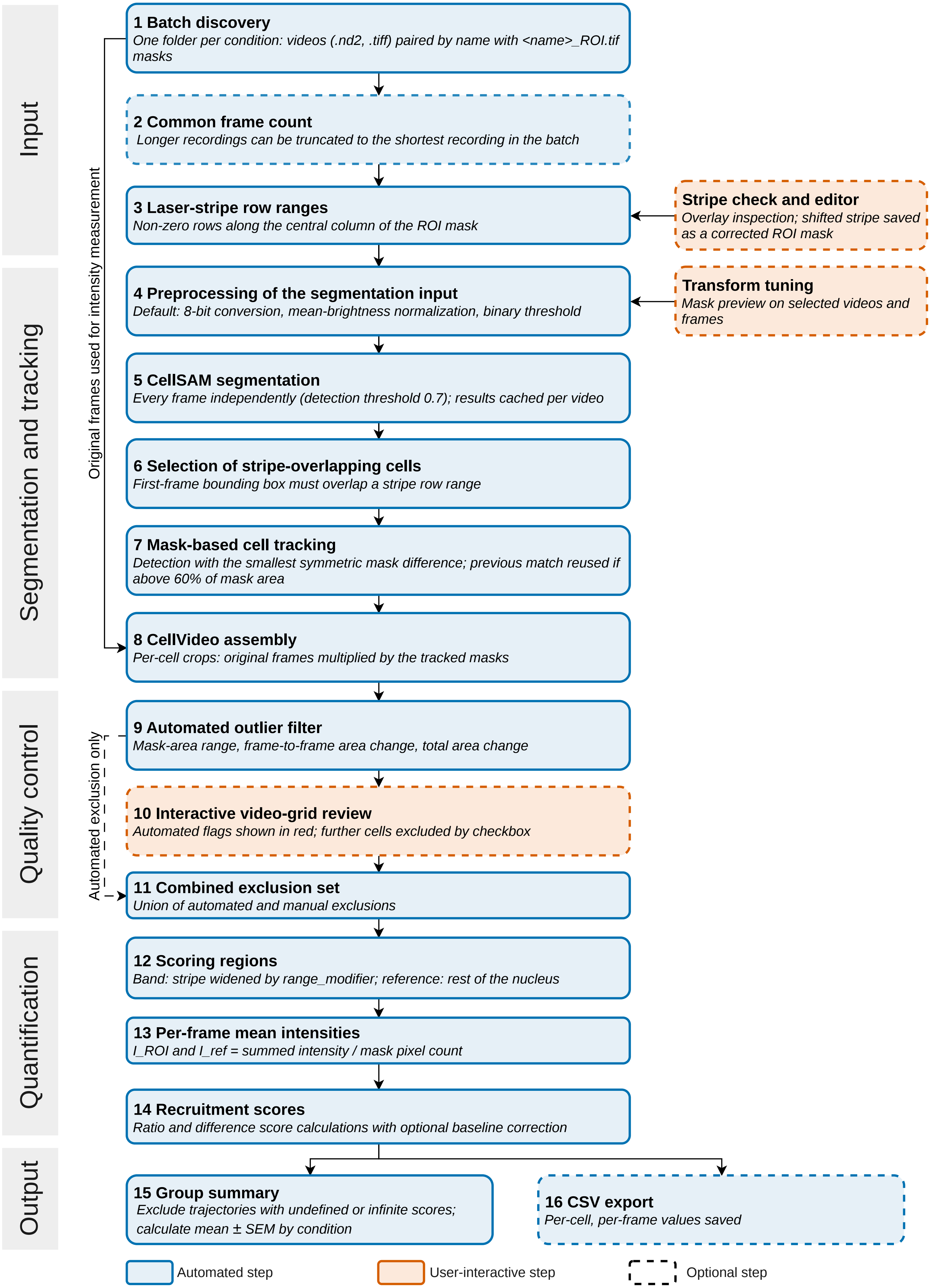
Input
Segmentation and tracking
Quality control
Quantification
Output
Original frames used for intensity measurement
Automated exclusion only
1 Batch discovery
One folder per condition: videos (.nd2, .tiff) paired by name with <name>_ROI.tif masks
2 Common frame count
Longer recordings can be truncated to the shortest recording in the batch
3 Laser-stripe row ranges
Non-zero rows along the central column of the ROI mask
Stripe check and editor
Overlay inspection; shifted stripe saved as a corrected ROI mask
4 Preprocessing of the segmentation input
Default: 8-bit conversion, mean-brightness normalization, binary threshold
Transform tuning
Mask preview on selected videos and frames
5 CellSAM segmentation
Every frame independently (detection threshold 0.7); results cached per video
6 Selection of stripe-overlapping cells
First-frame bounding box must overlap a stripe row range
7 Mask-based cell tracking
Detection with the smallest symmetric mask difference; previous match reused if above 60% of mask area
8 CellVideo assembly
Per-cell crops: original frames multiplied by the tracked masks
9 Automated outlier filter
Mask-area range, frame-to-frame area change, total area change
10 Interactive video-grid review
Automated flags shown in red; further cells excluded by checkbox
11 Combined exclusion set
Union of automated and manual exclusions
12 Scoring regions
Band: stripe widened by range_modifier; reference: rest of the nucleus
13 Per-frame mean intensities
I_ROI and I_ref = summed intensity / mask pixel count
14 Recruitment scores
Ratio and difference score calculations with optional baseline correction
15 Group summary
Exclude trajectories with undefined or infinite scores; calculate mean ± SEM by condition
16 CSV export
Per-cell, per-frame values saved
Automated step
User-interactive step
Optional step

**Supplementary Figure S2. Effect of additional manual filtering on automated XRCC1 recruitment analysis.**

Baseline-corrected fluorescence difference (Diff; A) and ratio (Ratio; B) recruitment curves for XRCC1 in siNT and siCyclin D1/D2/D3 cells. The plots were generated from the AUTO-derived dataset following additional manual filtering. Compared with the corresponding unfiltered AUTO results in Fig. 2A, this additional filtering did not substantially alter the overall recruitment patterns. Curves represent mean ± SEM and were additionally normalized by setting the highest mean recruitment value within each graph to 1.

**A**

mCherry-XRCC1 Diff (AUTO + additional filtering)

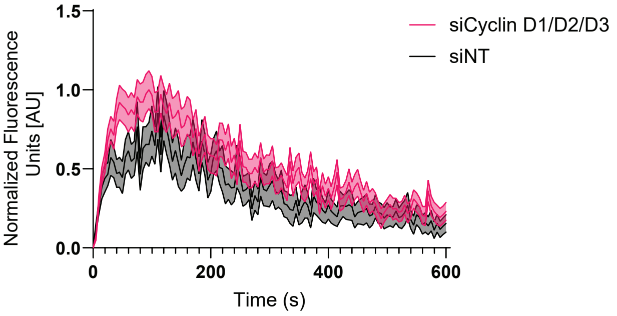


**B**

mCherry-XRCC1 Ratio (AUTO + additional filtering)

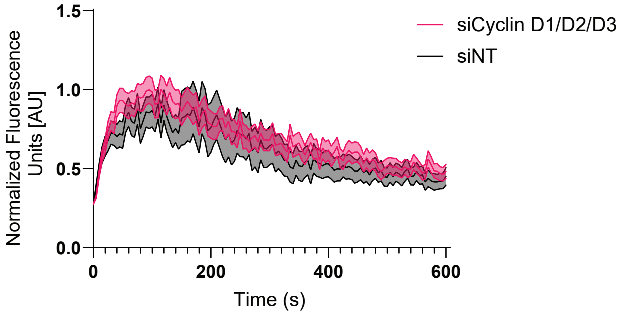


**Supplementary Figure S2**

**SUPPLEMENTARY TABLES**

**Supplementary Table S1.** DuckSeg analysis parameters used for the experimental datasets.

| Parameter | XRCC1 | NEIL3-C | NEIL3-N | MSH2 | MSH3 | EXO1b |
|---|---|---|---|---|---|---|
| Target mean brightness | 20 | 20 | 20 | 20 | 20 | 20 |
| Binary threshold | 20 | 20 | 20 | 20 | 20 | 20 |
| ROI width multiplier | 5 | 5 | 5 | 5 | 5 | 1 |
| Maximum nuclear area (pixels) | 7000 | 7500 | Not applied* | Not applied* | Not applied* | Not applied* |
| Interactive QC | No (AUTO); Yes (MANUAL) | No | Yes | Yes | Yes | Yes** |
| Acquisition interval (s) | 5 | 5 | 5 | 5 | 5 | 5 |
| Analyzed duration (s) | 600 | 600 | 495 | 605 | 605 | 725 |

NEIL3-C and NEIL3-N refer to C-terminally and N-terminally fluorescently tagged NEIL3, respectively. The ROI width multiplier corresponds to the range_modifier parameter.
* Maximum nuclear-area filtering was not applied.
** Interactive exclusion was used exclusively to retain S-phase cells.

**Supplementary Table S2.** Recruitment data underlying the manuscript figures.
Mean recruitment scores, standard deviations (SD), and numbers of contributing cells (N) at each time point for the experimental conditions presented in Figures 2 and 3 and Supplementary Figure S2. Values are provided following single-cell baseline correction but before the additional normalization applied in GraphPad Prism for graphical presentation. The number of contributing cells may vary between time points due to the inclusion of partially complete trajectories.
